# Exploring the Connection between Autophagy and Heat-Stress Tolerance in Drosophila melanogaster

**DOI:** 10.1101/2021.12.09.471892

**Authors:** Quentin Willot, Andre du Toit, Sholto de Wet, Elizabeth J. Huisamen, Ben Loos, John S. Terblanche

## Abstract

Mechanisms aimed at recovering from heat-induced damage are closely associated with the ability of ectotherms to survive exposition to stressful temperatures. Among these mechanisms the respective contribution of autophagy, a ubiquitous stress-responsive catabolic process, has more recently come to light. By increasing the turnover of cellular structures as well as the clearance of long-lived protein and protein aggregates, the induction of autophagy has been linked to increased tolerance to range of abiotic stressors in diverse ectothermic organisms. Since our understanding of the relationship between autophagy and heat-tolerance currently remains limited in insect models, we hypothesized that (1) heat-stress would cause an increase of autophagy in *Drosophila melanogaster* tissues and (2) rapamycin exposure would trigger a detectable autophagic response in flies and increase their heat-tolerance. In line with our hypothesis, we report that flies exposed to heat-stress present signs of protein aggregation and appears to trigger an autophagy-related homoeostatic response as a result. We further show that rapamycin feeding causes the systemic effect associated with TOR inhibition, induces autophagy at least locally in the fly gut, and increase the heat-stress tolerance of individuals. This points toward a likely substantial contribution of this autophagy to cope with stressful temperatures in insects.

## 1. Introduction

Without limited ability to regulate their internal body temperatures, terrestrial ectotherms’ physiology remains largely contingent on external temperatures and the degree of climatic variability they experience [1, 2]. They thus rely on a wide array of molecular-level processes to cope with fluctuation of environmental conditions and increase their resilience to physiological stress [3, 4]. Stirred by the projected issues for ectotherms related to climate change and exposure to more frequent episodes of climatic extremes [5, 6], the exploration of these molecular basis in invertebrates has received renewed attention [1, 7]. Most species are indeed capable of transient to permanent shifts in heat hardiness following prior exposure to stressful temperatures, a process commonly referred to as thermal acclimation, which increase the odds of survival to subsequent stress exposure [6, 8, 9]. Several adaptative and acclimation responses at the molecular-level have been correlated with enhanced tolerance to temperatures-induced stress, including, but not restricted to, the overexpression of heat-shock proteins (Hsps) [10–12], increased clearance of cellular protein aggregates [13, 14], modulation of cell-membrane phospholipids [15–17], or improved capacity for ion balance defence [18–20].

Within the molecular toolbox available to invertebrates to cope with environmental and energetic stressors, the contribution of autophagy recently enjoyed renewed attention [21–23]. Essentially, autophagy is a bulk lysosomal degradation pathway conserved in all eukaryotic cells aimed at clearing cytosolic components, targeting mainly organelles, long-lived proteins, and protein aggregates [24–26]. In doing so, autophagy serves a “house-keeping” function and eliminates old or damaged cellular components that may lead to the disruption of homeostasis [24], which in a context of temperature-induced stress is paramount for organisms [27, 28]. Three types of autophagic pathways can be recognized depending on how intracellular material reaches the lysosome [29]. Briefly, during macroautophagy, large cytoplasmic components or large protein aggregates can be captured within an autophagosome, the functional unit of autophagy, which then merges with the lysosomal system for digestion of its content [29]. Lysosomes can also directly capture smaller components and organelles through microautophagy [30]. Finally, denatured peptides can selectively be targeted to reach the lysosome via the assistance of Hsp70-family proteins, a process referred to as chaperone-mediated autophagy (CMA) [31]. In all cases, these autophagic pathways lead to the degradation of cellular targets. From a regulatory perspective, in *Drosophila melanogaster*, autophagy induction is under the control of the Target of Rapamycin (TOR) pathway [25, 32–35]. TOR inhibition in response to stress or exposure to rapamycin causes an cellular increase in autophagic flux and limits the synthesis of proteins that are not involved in the stress response [32]. This both directly mitigates the potential damages caused by abiotic stress while improving cellular metabolic robustness and survival by maximizing ATP and metabolite availability [36]. Modulation of autophagy, that relate to a more global responses from the TOR pathway, has therefore ultimately linked to enhanced survival to various stressors in animals, including insects, such as starvation, desiccation, toxics, heavy metals or oxidative damage [22, 23, 37–43]. In the specific case of heat-stress, it has been demonstrated that activation of the transcription factor HSF-1 which controls the cellular heat-shock response, and transcription of heat-inducible Hsps, also induces autophagy in multiple tissues in *Caenorhabditis elegans* [14]. This helps reduce the accumulation of denatured protein and increases chances of survival [14]. A link between rapamycin exposure and increased heat-tolerance in *D. melanogaster* has also been more recently suggested [44]. Taken together, this provides strong circumstantial evidence for the upregulation of autophagy to potentially be another pivotal aspect underpinning heat tolerance in insects.

Here, we thus aimed at exploring the potential mechanisms bridging rapamycin exposure, autophagy, heat-stress, and increased heat-tolerance in an insect model. We report that a rapamycin-enriched diet does induce the systemic effect associated with TOR inhibition in *D. melanogaster*, including impaired larval growth and increased formation of lysosomal acidic compartments in the fly gut. Second, we show that exposition to heat-stress further relates to accumulation of autophagy-related clearance proteins in *Drosophila* tissues, where a combination of rapamycin-exposure and heat-stress causes maximal abundance of GABARAPl1/2, Ref(2)P and Hsp70 in flies. Finally, we link rapamycin-exposure to enhanced tolerance to, and recovery from, heat stress. Putting autophagy within the context of heat-stress, we show that its induction can indeed play an important role nested within the cellular mechanisms enabling increased insect resilience to temperatures.

## 2. Methods

### 2.1 Fly Stocks and rearing

Wild *Drosophila* flies were bait-trapped in April 2019 around the Stellenbosch University campus, Western Cape, South Africa. Single female flies were isolated into 250 ml rearing flask supplied with 50ml of Bloomington cornmeal diet [45] and left to lay eggs for one day. Between 2 and 5 of the first emerging adults were used for species determination following the key of Markow TAO’Grady P [46]. Stocks were kept at a density of ~100 flies per flask, at a constant 25°C and a 12:12 light-dark cycle until experiments.

### 2.2 Rapamycin Treatment and heat-shock

For larval developmental assays, rapamycin (LC Laboratories) was dissolved in molecular biology grade absolute ethanol (Sigma-Alldricht) and added to the Bloomington cornmeal diet [45] at a concentration known to induce systemic effects in flies (50 or 200μM, [37]. For control (0μM), ethanol alone was added. For adults involved in dissection, microscopy, western blotting and heat-knockdown/recovery assays, eight days old flies were fed liquid food (5% yeast extract, 5% sucrose, 1% ethanol, 1% brilliant blue FCF) with the appropriate concentration of rapamycin (0, 50 or 200μM) for two days. Blue food commercial dye (Brilliant blue FCF, Robertson’s Blue Food Colouring) was used to visually confirm feeding.

For western blot analysis, rapamycin-treated adults were further either heat-shocked (37°C, 1h) or kept at 25°C to created 4 conditions within a full-factorial design: (1) Non rapamycin-exposed/non heat-shocked flies (control, rapa-/HS-), (2) rapamycin-exposed/non heat-shocked flies (rapa+/HS-), (3) rapamycin non-exposed/heat-shocked flies (rapa-/HS+), and (4) rapamycin-exposed/heat-shocked flies (rapa+/HS+). Flies were immediately flash-frozen after treatment and stored at −80°C for downstream analysis.

### 2.3 Developmental assays

Single 10-day old female flies were transferred into new vials containing the Bloomington cornmeal diet [45], supplemented with 0, 50 or 200μM rapamycin, and left to lay eggs for 24h. Development of larvae in the medium at 23°C was monitored daily, and number of pupations and eclosions recorded.

### 2.4 Heat-knockdown and recovery assays

For heat-knockdown experiments at 37°C, groups of 10 flies exposed to either 0, 50 or 200 μM rapamycin for 2 days were isolated into glass vials containing a moist cotton ball. Vials were then submerged in a programmable water bath (GD120 series, Grant Instrument Ltd) and the temperature inside the vials was monitored using 0.075 mm diameter thermocouples (Type T Thermocouple (Copper/Constantan), OMEGA Engineering, CT, USA) connected to a digital thermometer (Fluke 52-II Dual Input Digital Thermometer, WA, USA). Flies were exposed to a static 37°C heat-stress and checked every 15 minutes for loss of righting response, defined as the onset muscle coordination loss, when flies would fall on their back after a gentle vial shake and display total absence of movements. Between 110 and 130 flies were used for each experimental condition (0, 50 or 200 rapamycin-exposed) to extract knockdown curves at 37°C. For heat-knockdown experiments at 41°C, because the onset of muscle coordination would occur much faster, single flies were isolated into glass vials, exposed to 41°C, and monitored continuously. As soon as the loss of righting response occurred, each heat knocked-down fly was immediately placed back at room temperature (22°C) and monitored until recovery, defined as the time at which a fly could stand on its legs again, without stumbling (lack of coordinated movement) or falling over from external stimulation (gentle vial shaking). Between 45 and 51 flies were used to reconstruct knockdown/recovery curves at 41°C. Times to knockdown at 41°C and times to recovery were finally plotted against each other and the dataset screened for potential correlations.

### 2.5 Dissection, staining and confocal microscopy

To visualize the induction of autophagy in *D. melanogaster*, the gut of control or rapamycin-treated flies were dissected out in 4°C chilled PBS and then stained in a 10 μM Lysotracker Red:1μg/ml Hoechst solution (LysoTracker™ Red DND-99, Hoechst 33342 Solution, Thermofisher Scientific) for 3 minutes. Each preparation was washed three times with cold PBS, fixed overnight in 4% PFA at 4°C and mounted within the next 2 hours. Imaging was performed using a Zeiss LSM 780 ELYRA P.S.1 confocal microscope using a 20x objective. In each case, the midgut region between the proventriculus and Malpighian tubules were acquired. Laser power, digital gain and optical settings were kept constant between the images. Images were then binarized using a set threshold and pixel of high values quantified with the ImageJ software 1.50 [47]. The ratio of positive pixels per counting area was pooled for each individual and presented as an average made over between 7 and 13 individuals for each condition.

### 2.6 Western blotting

Following rapamycin/heat-shock treatments, whole fly bodies were collected, homogenized, and sonicated using a MixSonic (S-4000, ThermoFisher Scientific, Johannesburg, South Africa). The lysate was then centrifuged at 10 000 RPM for 20 min at 4°C (Spectrafuge, 24D, Labnet) to pellet nuclei and large cell fragments. The supernatant was collected into 1.5 mL Eppendorf tubes (#P2TUB003C-0001.5, Lasec, Cape Town, South Africa) and stored at −80°C. 15 mg of protein was mixed with Laemmli’s buffer (1 mL Tis-HCl (0.5 M; pH 6.8), 1.6 mL 10% sodium dodecyl sulphate (SDS, #MKCG7687, Sigma-Aldrich, Johannesburg, South Africa), 0.8 mL glycerol (#1045422, Merck, Johannesburg, South Africa) and 0.4 mL of 0.05% bromophenol blue (#1041675, Merck) in 3.8 mL dH2O) in a 2:1 ratio. Samples were subsequently boiled at 95°C for 5 min. Proteins were separated on a Fast Cast TGX Stain-Free gel (#456-8084, Bio-Rad, Johannesburg, South Africa) consisting of a 12% resolving and 4% stacking component, respectively (1610174, Bio-Rad) and transferred to a PVDF membrane (#170-84156, Bio-Rad) using the Trans-Blot Turbo (#170-4155, Bio-Rad).

Membranes were blocked for 2 h using 5% non-fat milk made up in 1x TBS-T (137 mM sodium chloride, 20 mM Tris, 0.1% Tween-20 at pH 7.6). Membranes were washed 3 times for 5 min using TBS-T and incubated with primary anti-bodies overnight at 4°C. Primary anti-bodies anti-Hsp70 (ab2787, Abcam, Germany), anti-Ref(2)P (ab178440, Abcam, Germany), and anti GABARAP1+ GABARAP2 (ab109364 Abcam, Germany) were used at a 1:5000 dilution in TBS-T. Membranes were washed three times with TBS-T for 5 min and incubated with peroxidase-linked anti mouse or rabbit IgG at a 1:10,000 dilution in TBS-T (#7076/#7074S, Cell Signaling, Germany) for one hour at room temperature. Finally, membranes were visualized using Clarity Western ECL substrate (#170-5061, Bio-Rad), prepared at a 1:1 ratio and acquired using the ChemiDoc MP System (Bio-Rad).

### 2.7 Statistical analysis

Statistical analyses were performed using GraphPad Prism 9.01. Kruskal-Wallis tests were followed by a Dunn’s multiple comparisons post hoc test. One-way ANOVAs were performed followed by a Fisher’s LSD post-hoc test. Significance for log-rank tests analyses of Kaplan-Meier survival and development curves, linear regressions, Kruskal-Wallis, and ANOVA tests were accepted at *p*<0.05.

## 3. Results

### 3.1 Rapamycin feeding delays Drosophila larval development

Delayed larval development because of TOR inhibition is a documented effect of rapamycin exposure in *D. melanogaster* (Zhang *et al.*, 2000). We thus performed larval growth assays to confirm that, in our experimental setup, a rapamycin-enriched diet would induce the expected systemic effects associated with TOR inhibition. Larvae were left to develop in Bloomington cornmeal diet (Lewis, 1960) supplemented with either 0 (control), 50 or 200 μM of Rapamycin. The number of pupations and imagoes hatching was recorded as a function time at 23°C. 50 μM rapamycin exposure significantly delayed the onset of first larval pupations by about 4 days and hatching by 6 days (Fig.3; Log-ranked test, p<0.001). Survival from first instar to imago was estimated at 98% in control conditions (N=453) and dropped to 36% with 50 μM supplemented rapamycin food (N=129). No pupation events were observed at 200 μM (all larvae remained either in second or third instars until mortality reached 100% after 20 days).

### 3.2. *Rapamycin feeding induces modulation of autophagy in the* Drosophila *midgut*

Adult flies were fed control or rapamycin supplemented food at 50 and 200μM. Midguts of flies were dissected and stained with Lysotracker prior to imaging. On average, there was a significant increase in the number of Lysotracker-positive compartments observed in the midgut of flies after rapamycin treatment, suggestive of drug uptake in the digestive track and local autophagy induction (Fig.2A, B). Quantification of the average number of pixels above intensity threshold in midgut areas of flies revealed of a dose-dependent response as a function of the rapamycin concentration in their food (Fig. 2C).

**Figure 1.**
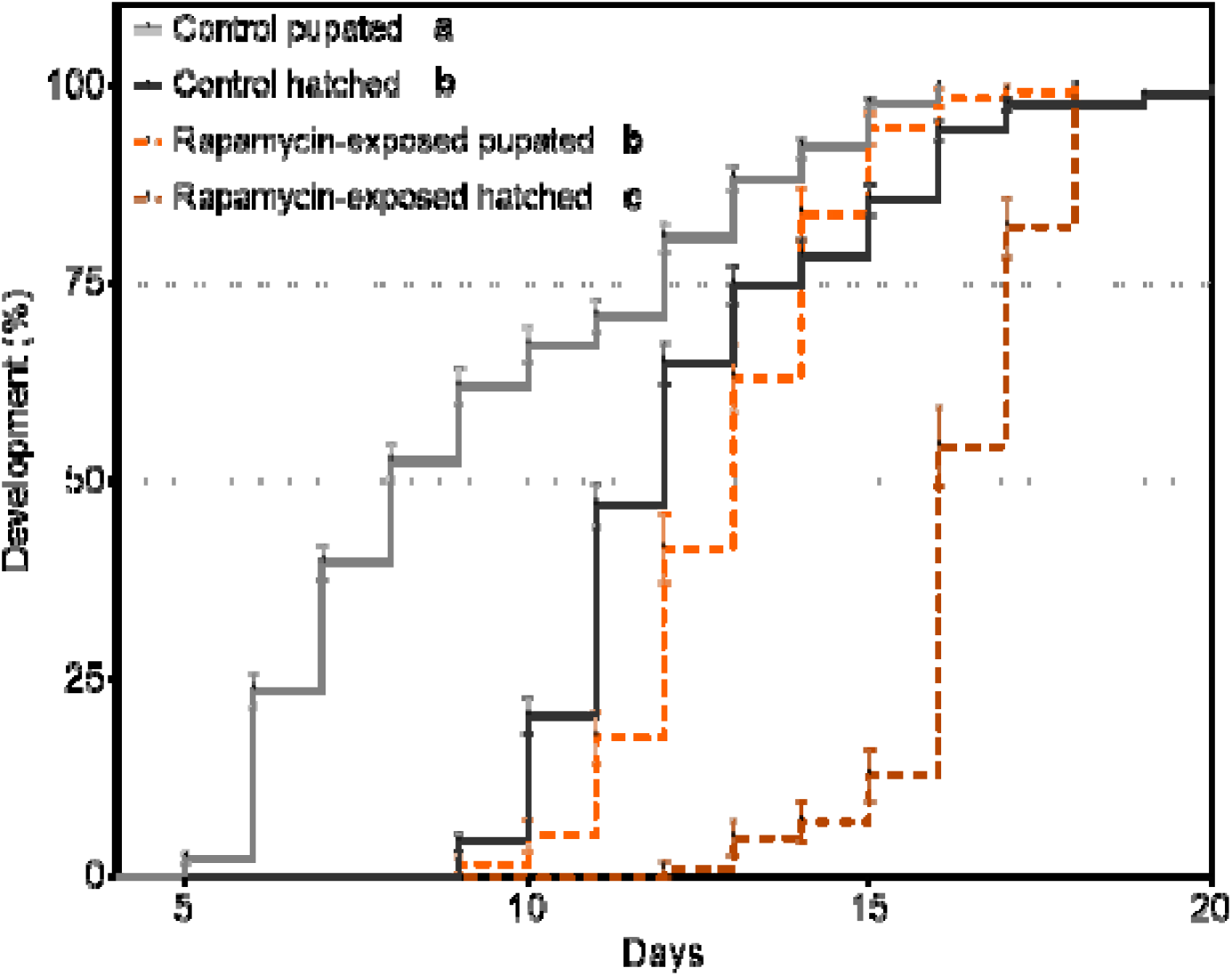
Number of *Drosophila melanogaster* pupation and hatching events as a function of time at 23°C, for larvae fed either control (0 μM) or a 50 μM rapamycin supplemented diet. Rapamycin exposure delayed both the onset of pupation and hatching by several days. Different lowercase letters indicate significant differences between treatments (Log-rank tests, P<0.001).

**Figure 2.**
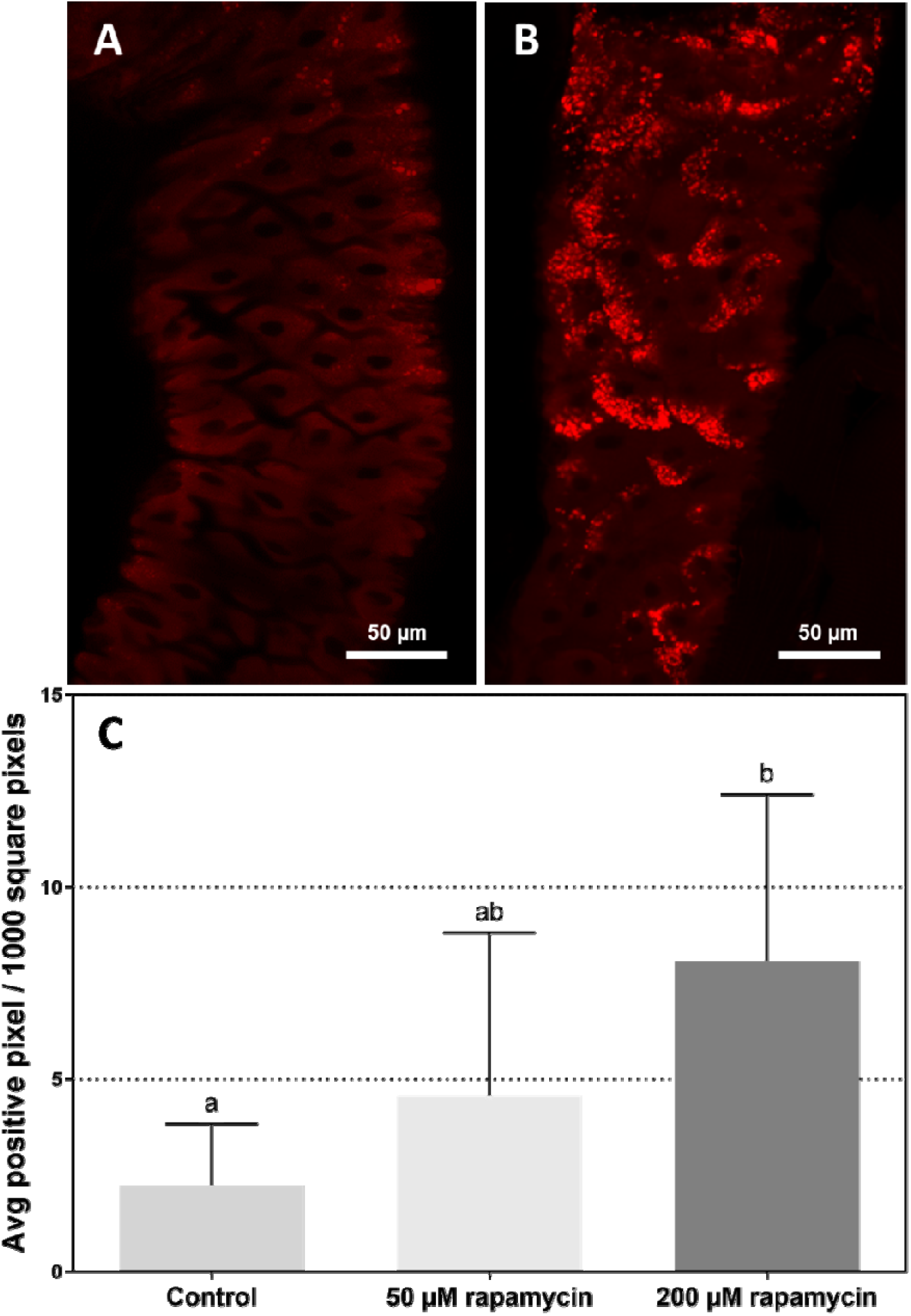
Representative confocal fluorescence images of adult *Drosophila* midguts stained with LysoTracker Red (scale bar is 50 μM). **A.** Midgut of control flies. **B.** Midgut of rapamycin treated flies (200 μM). **C.** Average number of pixels above intensity threshold over midgut images of 7 to 13 individuals per conditions. Different lowercase letters indicate significant differences between treatments (Kruskal-Wallis one-way analysis of variance followed by Dunn’s multiple comparisons post hoc test, p<0.01).

**Figure 3.**
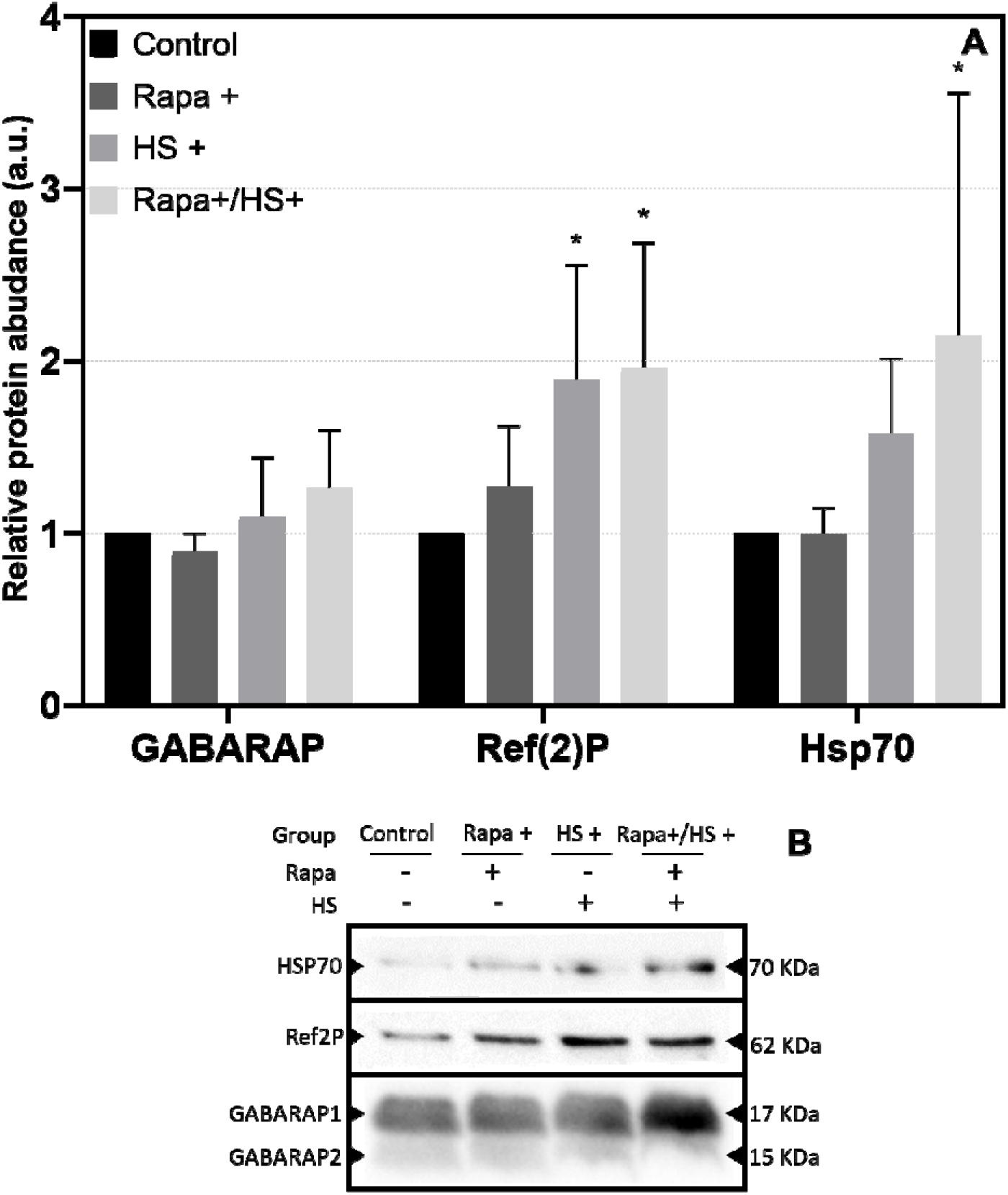
Western blots analysis (N=4) following 200 μM rapamycin exposure of fly tissues directed against GABARAP, Ref(2)P and Hsp70. **A.** Relative abundance to control of GABARAPL1/2, Ref(2)P and Hsp70-family proteins following rapamycin exposure and/or heat-shock. GABARAP reached near-statistical significance following heat-stress in rapamycin treated conditions (*p*=0.053). Ref(2)P levels were significantly up-regulated following both heat-stress and heat-stress/rapamycin exposure conditions (*p*<0.05). Finally, Hsp70 levels were not impacted by rapamycin exposure alone, but increased following heat-shock to reach peak level through a combination of rapamycin exposure and heat-shock (*p*<0.05). Asterix represent significant differences relative to control (One-way analysis of variance followed by a Fisher’s LSD post-hoc test, *p*<0.05). **B.** Representative western blots are shown.

### 3.3. Exposition to heat-stress and rapamycin modulates autophagy in fly tissues

A qualitative confirmation of engagement of the autophagosome/lysosome system was performed on fly issues through western blot analysis (Fig 2A, N=4). Flies were either kept on control conditions (rapa-/HS-) or exposed to rapamycin and/or heat-shock (rapa+/HS-, rapa-/HS+, rapa+/HS+), and blotting was performed against: (1) GABARAPL1/2, which are Atg8 protein family members involved in autophagosome formation [48], (2) Ref(2)P, the *Drosophila melanogaster* ortholog of p62/sequestosome-1, involved in directing aggregated protein cargoes towards autophagic/proteasomal degradation pathways by acting as receptor [49, 50] and (3) HSP70-family proteins that are hallmark of proteotoxic stress and involved both in the peptide-refolding machinery and CMA [31, 51]. GABARAPL1/2 levels reached near-statistical significance following heat-stress in rapamycin-treated conditions (*p*=0.053) (Fig. 3A). Ref(2)P abundance as compared to control was significantly increased following heat-stress and the combination of heat-stress and rapamycin exposure (*p*<0.005). Finally, the abundance of Hsp70 was not impacted by rapamycin alone but increased following heat shock and the combination of rapamycin and heat-shock as compared to control (*p*<0.05; Fig. 3A). Representative western blots are shown (Fig. 3B).

### 3.4. Rapamycin treatment delays time to knockdown in flies at 37°C

Eight-day old adult flies were fed a rapamycin-enriched diet at either 0 (control), 50 or 200 μM for two days, and then subsequently exposed to a static 37°C stress to induce a time-dependent heat knockdown. The number of knocked down flies as a function of time was recorded every 15 minutes (Fig. 4). Rapamycin exposure delayed the time for flies to enter heat knockdown (Log-ranked test, p<0.001). 50 μM rapamycin exposure in food was already sufficient to induce a plateau response.

**Figure 4.**
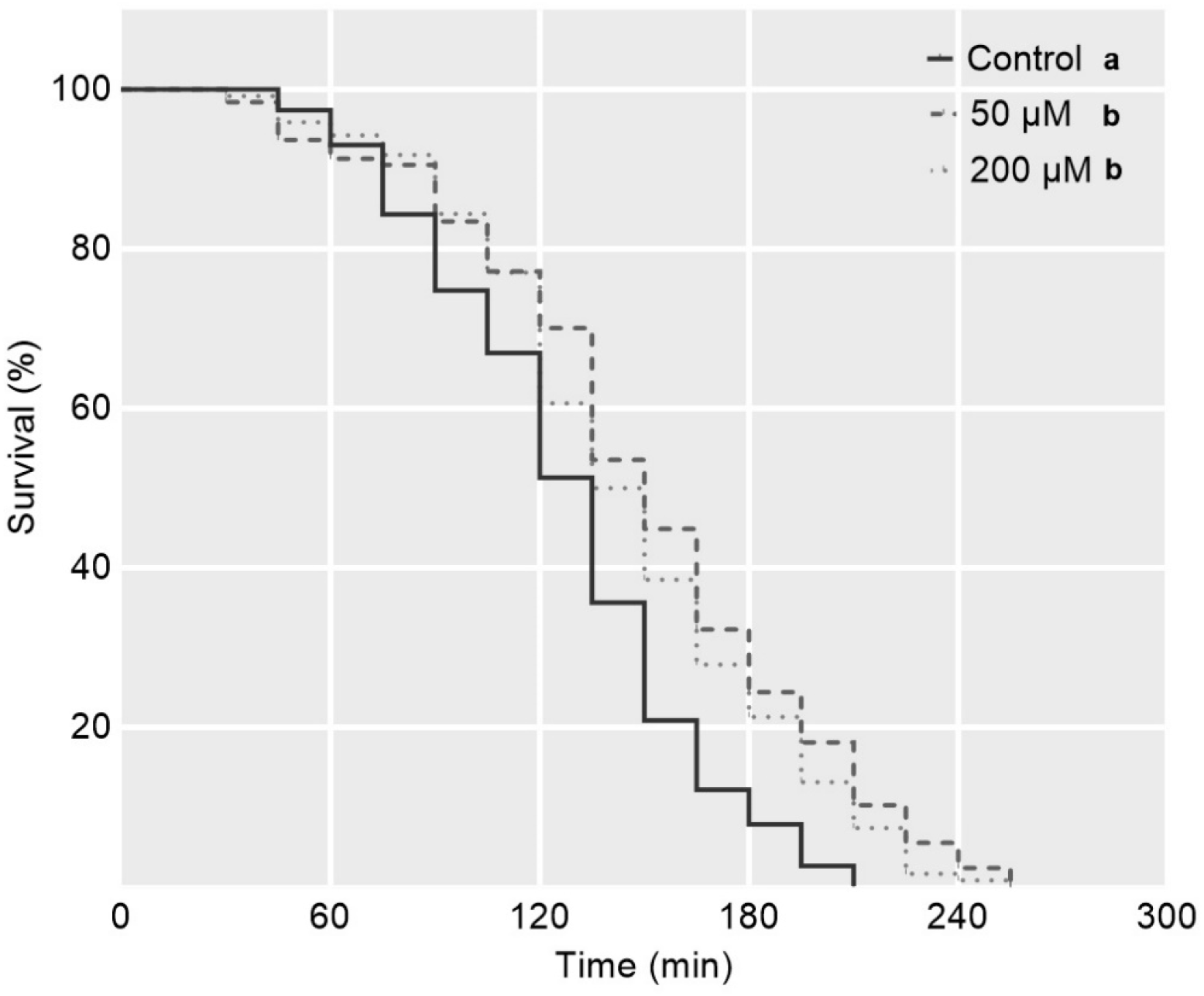
Survival of *Drosophila melanogaster* flies that were exposed to a rapamycin-enriched diet to a 37°C static heat stress (N=115 to 130 per condition). Rapamycin exposure significantly delayed time-to heat knockdown in flies, and 50 μM exposure was sufficient to induce a plateau response. Different lowercase letters indicate significant differences between treatments (Log-rank tests, P<0.001).

### 3.5. Rapamycin treatment decreases time to recovery post knockdown

Eight-day old adult flies were fed a rapamycin-enriched diet at either 0 (control), 50 or 200 μM for two days and subsequently exposed to a static 41°C stress to induce a fast heat-knockdown. Flies were monitored continuously and put back at room temperature (23°C) for recovery after the loss of muscle coordination. The impact of rapamycin exposure on heat knockdown at 41°C and its linked recoveries are presented in Fig. 5. For heat knockdown at 41°C, no effect of rapamycin exposure was observed (Fig. 5A). However, rapamycin exposure reduced the time to recovery post knockdown in flies (Fig. 5B, Log-ranked test, p<0.001). Finally, for each individual fly, single times to knockdown at 41°C were plotted against single times to recovery, and significant regressions were screened both as a function of rapamycin exposure (Fig. S1A) and on the global dataset (Fig. S1B; Table S1). Only a weak correlation between time to heat knockdown and heat recovery was found for flies fed with 50 μM rapamycin (R^2^=0.22, p <0.001, Fig. S1A) and on the global dataset (R^2^=0.05, p<0.01; Fig. S1B). No significant correlations were recorded for other conditions (Table S1).

**Figure 5.**
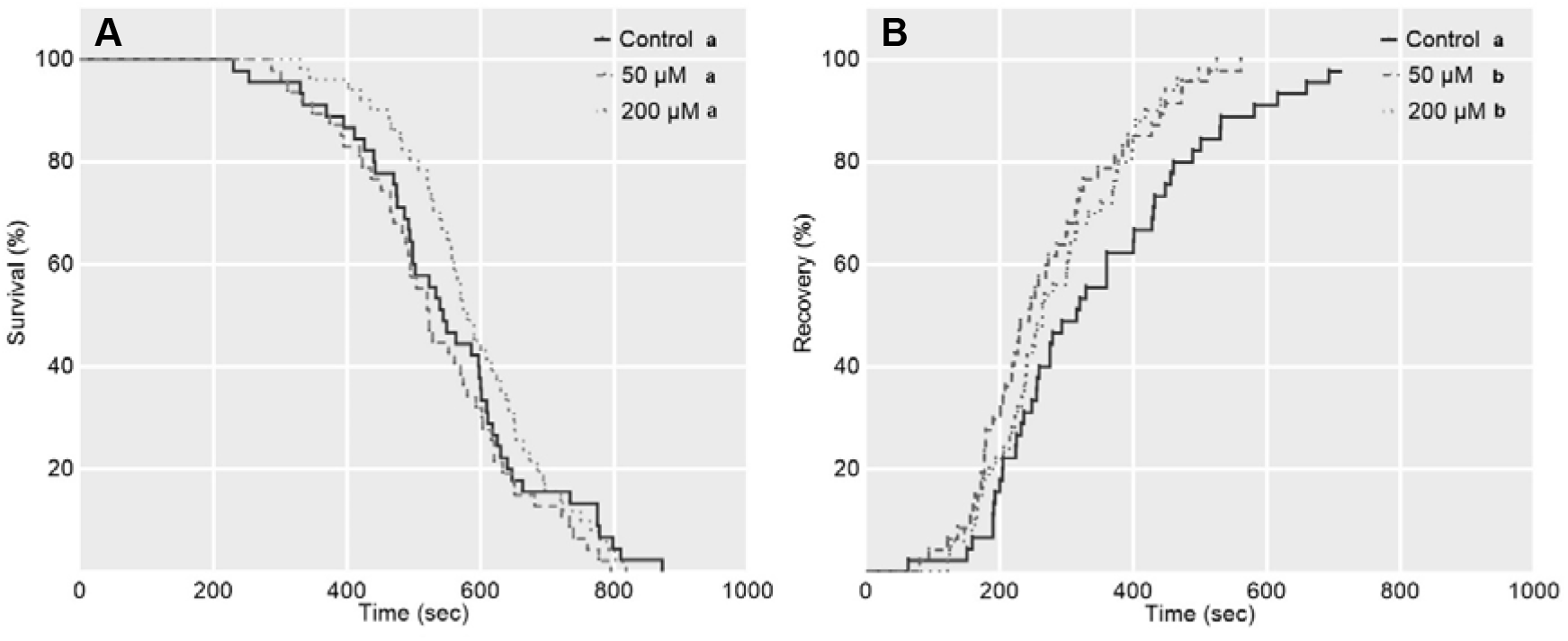
knockdown and recovery curves of *D. melanogaster* flies that were exposed to a rapamycin-enriched diet as a function of time at 41°C. **Fig. 5A.** Rapamycin exposure did not delay time to heat knockdown in flies at 41°C. **Fig. 5B.** Rapamycin exposure decreased time to recovery from heat knockdown of flies. Different lowercase letters indicate significant differences between treatments (Log-rank tests, *p*<0.001).

## 4. Discussion

A number of markers to track autophagy have been validated in *Drosophila* so far, among which the Atg-8 family protein can potentially track all stages of the autophagy pathway by providing key information on autophagosome abundance [48, 52, 53] and the accumulation of ref(2)p which directs ubiquitinated proteins/proteins-aggregates towards the proteasome and/or selective autophagy [49, 50, 52, 53]. While these two markers target complementary, independent functions within the scope of autophagic activities [54], these are limited in obtaining reliable overviews of the autophagic rate (or flux) because of the inherent dynamic nature of this process [52, 55]. For example, the number of autophagosomes in tissues can increase either due to enhanced synthesis or due to accumulation brought by lysosomal dysfunction [56], and since autophagosome synthesis can be balanced by increased degradation, enhanced autophagic rate might result in constant Atg-8 level estimates at a snapshot despite the autophagy flux being upregulated towards a new steady state flux [52, 53, 57]. To circumvent these problems, the use of lysosomal inhibitors such bafilomycin or chloroquine are often used *in vitro* [52, 53, 55, 57]. However, these are poorly compatible with studies involving intact larvae or adult flies because of dire secondary effects that the general inhibition of lysosomal degradation has on the physiology of whole organisms [52]. Likewise, in live *D. melanogaster* larvae and flies, inhibition of the TOR signalling pathway by rapamycin exposure bears global effects on metabolism via e.g., increased autophagic turnover, and limited *de-novo* protein synthesis. These effects have been documented from hindered larval growth and delayed imago eclosions [33], to increased lysotracker-positive punctate in the fly gut which is indicative of local autophagy induction, and increased survival to starvation and paraquat [37]. Thus, to link autophagy induction with enhanced survival to heat-stress within the boundaries of our own experimental system, it was imperative to first confirm that a rapamycin-rich diet would indeed result in the expected effects associated with the drug-exposure in both larvae and adults. Consistent with previous findings, we confirm that exposure to rapamycin does indeed significantly delay *D. melanogaster* larval development (Fig. 1), and that a 2-days exposure to rapamycin-enriched diet causes a dose-effect response in the number of lysotracker-positive punctate in the fly midgut (Fig. 2). Taken together, this provides strong circumstantial evidence that rapamycin-feeding as performed during our treatments indeed causes the documented effects associated with TOR inhibition and promotes autophagy, at the very least, locally in the fly gut.

Based on these results, we attempted to detect whether rapamycin-feeding would cause autophagy induction at the systemic level in the fly, by tracking autophagic markers in tissues through western-blotting (Fig. 3). We could not detect a statistical significant effect of rapamycin exposure alone on GABARAPL1/2 levels (Atg-8 family member) [48] nor on Ref(2)P abundance (Fig. 3A) in fly tissues. One likely potential explanation for this could be found in the limited temporal resolution offered by our experimental design, as flies were exposed for 2-days to the drug prior to experiments. Markers such has Ref(2)P have been shown to decrease rapidly following autophagy induction but restored during such prolonged stress durations [58], and it could well be that a 2-days rapamycin exposure period might simply fail to track changes in marker abundance within their relevant timeframe. However, the near-statistical increased of GABARAPL1/2 abundance (*p*=0.053), and the increased abundance of Ref(2)P and Hsp70 (*p*<0.05) in either heat-stressed and/or heat-stressed/rapamycin exposed flies offer interesting perspectives (Fig. 3). First, increased GABARAPL1/2 levels following heat-stress and especially following heat-stress and rapamycin exposure could potentially reflect autophagosome accumulation through moderate autophagy induction [48]. Second, because Ref(2)p binds to ubiquitin-positive aggregates before degradation through the autophagic/proteasome pathway [49, 50], its accumulation can either reflect reduced degradation rate of protein aggregates or, more likely under the circumstances, an expected increase in protein aggregation associated with heat-stress [59–61]. Finally, and while we initially used well established Hsp70-family proteins as an internal marker for successful heat-stress exposure in flies [10], its peaking levels through the combination of heat-stress and rapamycin exposure could point toward an attempt for chaperone-mediated autophagy to recycle denatured peptides before they precipitate in toxic aggregates [31, 62]. While the full extent of these interpretations remains merely speculative without further testing, and while we could not statistically confirm autophagy induction in fly tissues following rapamycin-feeding alone probably because of the practical time-resolution limitation that a 2-day drug exposure offers, our results nonetheless suggest heat-stress does induce an expected proteotoxic stress in fly tissues and, to a degree, that an autophagy-related homoeostatic response might be triggered as a result thereof.

Finally, we show that rapamycin-exposure increases heat-tolerance in flies (Fig. 4). Albeit to the best of our knowledge no direct link between rapamycin exposure, autophagy induction, and heat-stress survival had been made in insect models prior to this work, this echoes with previous findings involving rapamycin feeding to cause a slight but significant increase of the critical thermal limits of *D. melanogaster* flies [44], and the overall involvement of autophagy in surviving an array of deleterious conditions in ectotherms [21–23, 37–43, 63, 64]. Solid support for a role of autophagy in mitigating heat-induced damages and increasing survival in invertebrates is also recorded from *Caenorhabditis elegans* where a causal link between autophagy induction, heat-acclimation and increased upper heat tolerance was made [14]. In perspective, we thus first show in *D. melanogaster* flies that 50 μM rapamycin exposure is already sufficient to induce a plateau-response stress survival under a 37°C stress exposure (Fig. 4, such delayed heat-knockdown was not observed in experiments conducted at 41°C which likely underpins that knockdown occurred too fast in this condition for us to detect a statistically significant response, Fig. 5A). Second, we were able to detect a benefit of rapamycin exposure in reducing recovery time post heat-knockdown (Fig. 5B). Although we provide evidence of a good correlation between these phenotypic effects and autophagy induction in response to rapamycin and heat-stress exposure in flies, we can only theorize on the responsible underlying molecular mechanisms causing the observed heat-tolerance increase. TOR inhibition and autophagy induction have been linked with a number of non-mutually exclusive processes related to cellular metabolic resilience and repair mechanisms, including increased clearance of toxic protein aggregates [14, 65], increase clearance of old and/or denatured peptides [62], optimization energy production and cellular metabolic state to maximize survival [36] and finally, recycling of damaged/surplus organelles such as mitochondria [23, 24]. Several of these could be jointly involved in the observed phenotypic increase in heat-tolerance in flies, and further work would be necessary to test and disentangle the respective contribution of each of them within the framework of autophagy induction.

In conclusion, our results show that both rapamycin and heat-stress support degrees of autophagic induction in *D. melanogaster*, and that rapamycin exposure correlates with an increase in heat-tolerance and speed of recovery post-knockdown in adults. This highlights the induction of autophagy as potentially key mechanism to increase resilience to temperatures in insects. A deeper exploration of autophagy’s role in relation to other stress responsive pathways remains however necessary and could provide important perspectives on the broader integrated cellular stress-response to temperatures in ectotherms.

## Supporting information

Extra tables and figures

## Authors’ contribution

Q.W., B.L. and J.S.T. conceived and planned the study. Q.W. and E.H. collected samples. Q.W., A.dT and S.dW performed lab experiments, Q.W. and S.dW analysed the data. All authors contributed to drafting the article, approved the final published version, and agree to be held accountable for all aspects of the work

## Acknowledgment

We thank Lize Engelbrecht and the team from the Central Analytical Facility of Stellenbosch University for their help with confocal imagery analysis.

## Preprint

A earlier version of this manuscript can be found in bioRxiv [66].

## Competing interests

We declare we have no competing interests.

## Funding

This work was supported by a Claude Leon Foundation post-doctoral fellowship (to Q.W.) and a running cost funding (to J.T.) from the DSI-NRF Center of Excellence for Invasion Biology.

## Notes

### Competing Interest Statement

The authors have declared no competing interest.

### Summary of Updates

Extra material

