## Supplementary material for "Exploring the Connection between Autophagy and Heat-Stress Tolerance in Drosophila melanogaster": Extra tables and figures

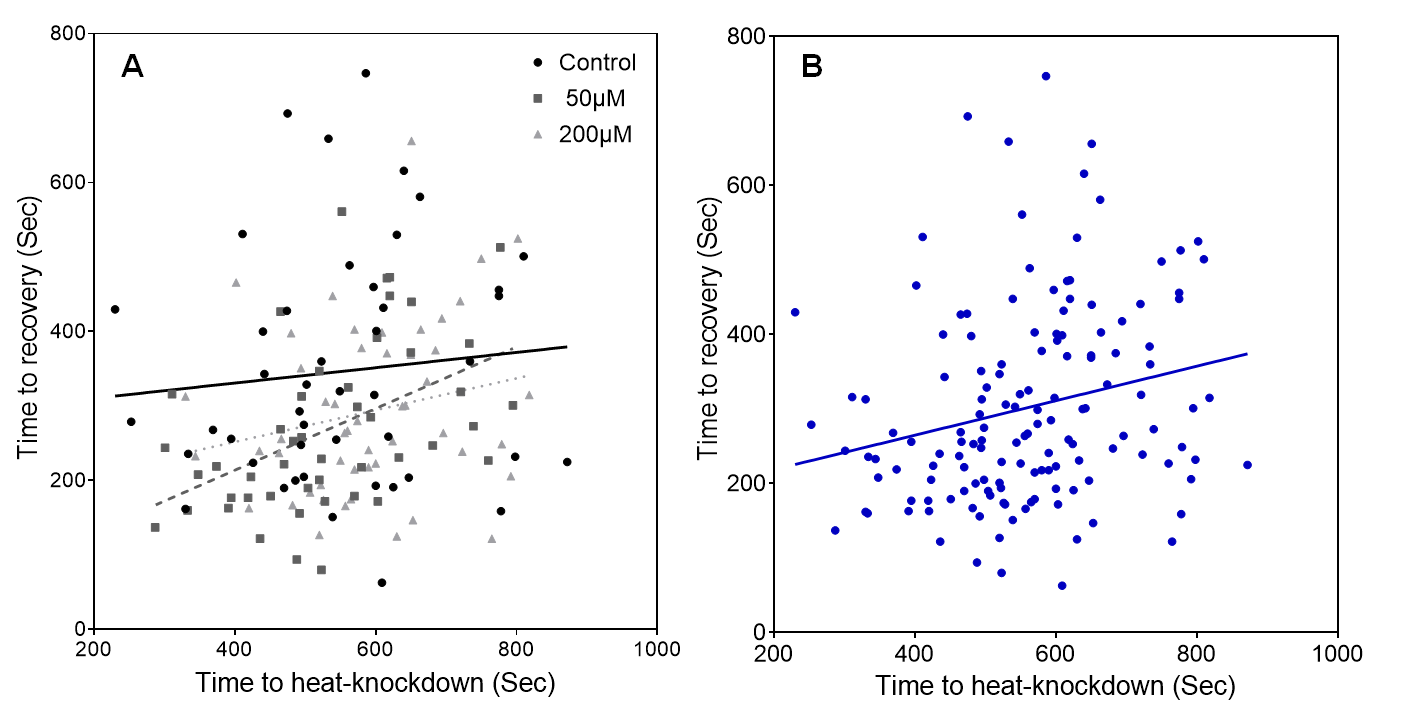

**Figure S1.** Time to recovery as a function of time to heat-knockdown at 41°C in individual flies**. Fig. S1A**: Linear regression for individuals pooled as a function of treatment. No significant correlation was observed, with exception of a weak linear relationship in 50 µM rapamycin fed flies (R²=0.22, p<0.001; Table 1). **Fig. S1 B**: regression on the global dataset, a minor but statistically significant linear correlation was present (R²= 0.05, *p*<0.01; Table 1).

**Table S1.** Details of the linear regressions calculated for time to recovery as a function of time to heat-knockdown at 41°C in individual flies. A weak but statistically significant relationship was observed for flies fed rapamycin at 50 µM and on the global dataset.

| **Condition** | **Equation** | **R²** | ***p* value** |
| --- | --- | --- | --- |
| Rapamycin 0 µM (Control) | Y = 0,1028*X + 289,9 | 0.008 | 0.54 |
| Rapamycin 50 µM | Y = 0,4141*X + 48,21 | 0.220 | 9*10^-4^ |
| Rapamycin 200 µM | Y = 0,2135*X + 166,5 | 0.044 | 0.13 |
| Global dataset | Y = 0,2315*X + 172,3 | 0.051 | 6.3*10^-3^ |
